# Self-organization of a doubly asynchronous irregular network state for spikes and bursts

**DOI:** 10.1101/2021.03.29.437548

**Authors:** Filip Vercruysse, Richard Naud, Henning Sprekeler

## Abstract

Cortical pyramidal cells (PCs) have a specialized dendritic mechanism for the generation of bursts, suggesting that these events play a special role in cortical information processing. *In vivo*, bursts occur at a low, but consistent rate. Theory suggests that this network state increases the amount of information they convey. However, because burst activity relies on a threshold mechanism, it is rather sensitive to dendritic input levels. In spiking network models, network states in which bursts occur rarely are therefore typically not robust, but require fine-tuning. Here, we show that this issue can be solved by a homeostatic inhibitory plasticity rule in dendrite-targeting interneurons that is consistent with experimental data. The suggested learning rule can be combined with other forms of inhibitory plasticity to self-organize a network state in which both spikes and bursts occur asynchronously and irregularly at low rate. Finally, we show that this network state creates the network conditions for a recently suggested multiplexed code and thereby indeed increases the amount of information encoded in bursts.

**Author summary:** The language of the brain consists of sequences of action potentials. These sequences often contain bursts, short “words” consisting of a few action potentials in rapid succession. Bursts appear to play a special role in the brain. They indicate whether a stimulus was perceived or missed, and they are very effective drivers of synaptic plasticity, the neural substrate of learning. Bursts occur rarely, but consistently, a condition that is thought to maximize the amount of information they can communicate. In our article, we argue that this condition is far from self-evident, but requires very special circumstances. We show that these circumstances can be reached by homeostatic inhibitory plasticity in certain inhibitory neuron types. This may sound complicated, but basically works just like a thermostat. When bursts occur too often, inhibition goes up and suppresses them. When they are too rare, inhibition goes down and thereby increases their number. In computer simulations, we show that this simple mechanism can create circumstances akin to those in the brain, and indeed allows bursts to convey information effectively. Whether this mechanism is indeed used by the brain now remains to be tested by our experimental colleagues.

## Introduction

Cortical activity consists of irregular sequences of spikes^1^, interspersed with bursts of several action potentials in quick succession^2,3^. Many cells in the nervous system have specialized cellular mechanisms for the generation of bursts^4–6^, suggesting that they play a special role in cortical information processing. Burst activity has been associated with a variety of computational and cognitive functions, including the conscious detection of stimuli^7^, the reliable transmission of information^8^ and the induction of synaptic plasticity^9^.

Most burst generating mechanisms rely on nonlinear membrane dynamics and are triggered by specific input conditions ^4–6^. In pyramidal cells, bursts can be generated by a coincidence of back-propagating actions potentials and synaptic input to the apical dendrite^4^. This associative mechanism could underlie the integration of external sensory signals — reaching the peri-somatic domain — and internally generated signals^10^ such as predictions^11,12^ or errors^13–16^, which reach the apical dendrite in superficial cortical layers. Based on the observation that different information streams target different compartments that in turn generate distinct spike patterns, it was recently suggested that both information streams could be conveyed simultaneously by means of a multiplexed neural code^17^. Such a multiplexing could be exploited, e.g., to route feedforward and feedback information in hierarchical networks^13,17,18^.

For bursts to convey information effectively, they need to occur rarely, but consistently^17^. Neural recordings suggest that this is indeed the case^2,3,7^. Given that bursts are triggered by nonlinear threshold-like processes, however, this state is not easily reached^1^. If dendritic input to PCs is too low, bursts will be absent entirely. If it is too high, bursts will be the predominant form of activity. Both conditions limit the amount of information bursts can transfer. This suggests that neurons should homeostatically regulate the amount of bursts they emit, by controlling dendritic excitability or the amount of dendritic input they receive.

A potential candidate for such a homeostatic control of burst activity is dendritic inhibition. Apical dendrites of cortical PCs receive inhibition from distinct inhibitory interneuron classes ^19,20^, including somatostatin-expressing (SOM) Martinotti cells^21^. SOM interneurons could be highly effective homeostatic controllers of burst activity, because the dendritic plateau potentials that underlie burst generation in PCs are very sensitive to inhibition^4,22,23^. Yet, this high sensitivity asks for dendritic inhibition that is finely tuned to the level of dendritic excitation, i.e., dendritic inhibition should be adaptive. Such a mechanism of preserving a suitable level of dendritic inhibition has been theorized to be essential for dendrites to participate in the coordination of synaptic plasticity ^13^.

Here, we use a computational network model to show that such a homeostatic control could be achieved by a simple form of dendritic inhibitory plasticity. We show that this plasticity can be readily combined with other forms of inhibitory plasticity that control cellular activity levels overall^24^. In recurrent spiking networks, the combination of these two forms of inhibitory plasticity can establish a doubly irregular state, in which both spikes and bursts occur irregularly at a consistent, but low rate. Finally, we show that inhibitory plasticity can self-organize dendritic input levels such that a multiplexing of feedforward and feedback input^17^ is more robustly preserved.

## Results

In L5 PCs, bursts occur at a low, but consistent rate ^2,3,7^ and are thought to originate from active dendritic processes ^4^. We hypothesized that this is the result of a homeostatic control of burst firing, mediated by plasticity of inhibitory synapses onto apical dendrites. But which inhibitory plasticity rules could achieve such a control and what would be the consequences at the network level?

To address these questions, we used a current-based spiking network model consisting of excitatory PCs and inhibitory interneurons. To model the burst mechanism of L5 PCs, PCs were described by a simplified two-compartmental model^17,25^. In short, the somatic and dendritic compartment of the PCs are each modelled by a set of differential equations of the adaptive integrate-and-fire type, with a nonlinearity in the dendritic compartment that allows an active generation of dendritic spikes. The two compartments receive bottom-up and top-down input, respectively (Fig 1A) and communicate by passive and active propagation. This model faithfully predicts spike timing of PCs in response to electrical stimulation^25^ and reproduces the qualitative features of burst activity when L5 PCs are injected with somatic and dendritic input Fig 1B; ^4,17,25^. Inhibitory interneurons were described by an integrate-and-fire model. For clarity, we gradually increase the complexity of the network from an uncoupled population of PCs to a feedforward network with inhibition and, finally, a recurrent network with two interneuron classes, representing dendrite-targeting SOM interneurons and soma-targeting parvalbumin-positive (PV) interneurons. The parameters of the interneuron model were adjusted to reflect the properties of these cell classes, specifically the presence and absence of spike-frequency adaptation in SOM and PV neurons, respectively (see Methods).

**Figure 1.**
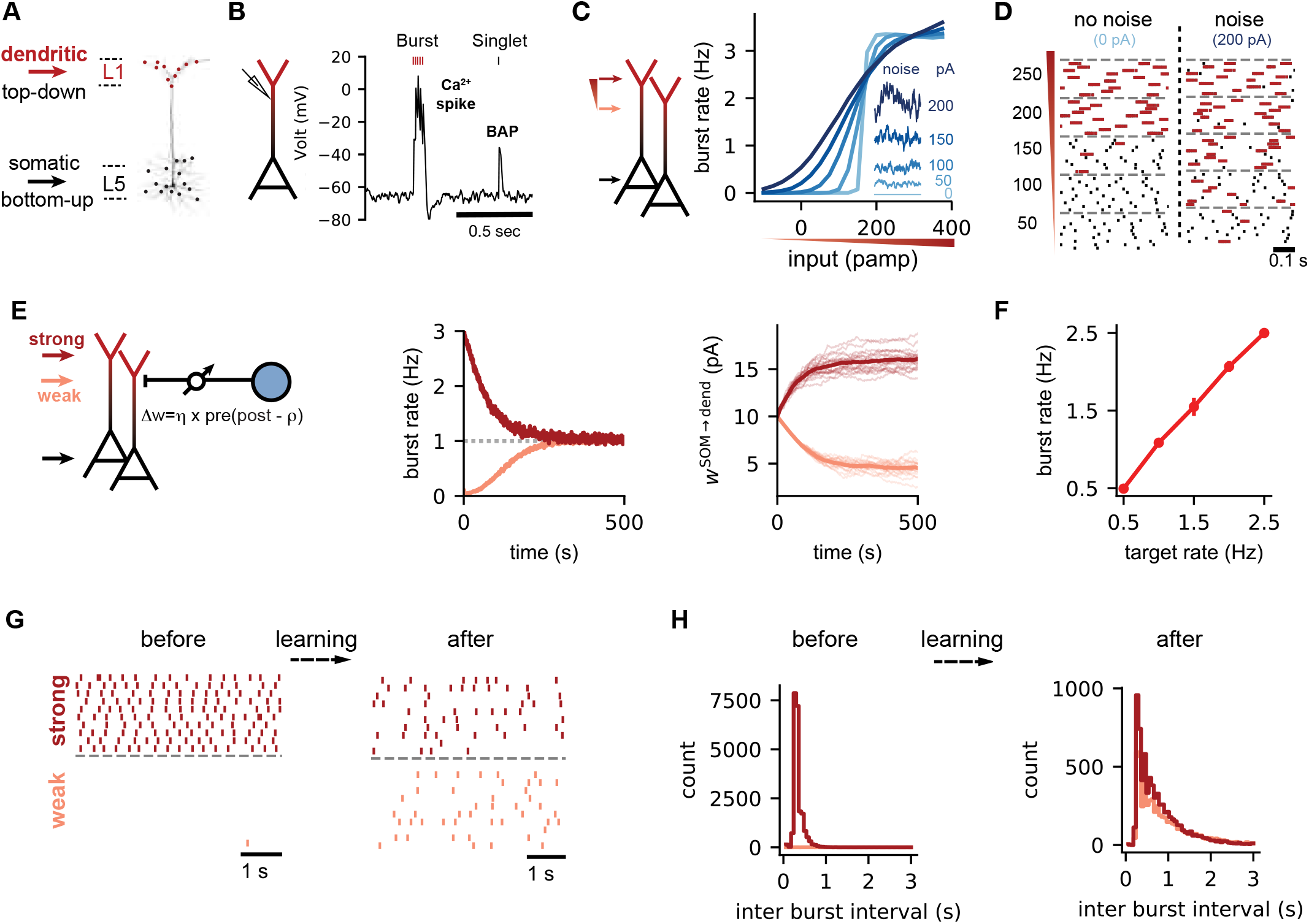
Control of the burst rate by homeostatic inhibitory plasticity. **(A)** Anatomy of layer 5 pyramidal neurons (L5 PCs). Sensory bottom-up inputs innervate the perisomatic region (black) while long range top-down connections target the distal dendrites (red). **(B)** Simulated dendritic voltage in a two-compartmental model of L5 PCs. A single somatic spike back-propagates into the distal dendrites. The coincidence of a back-propagating action potential (BAP) with sufficient synaptic input leads to sustained depolarization of the dendrites (calcium spike) and burst activity in the soma^4,17,25^. **(C)** L5 PCs are stimulated with varying degrees of dendritic input, characterised by an Ornstein-Uhlenbeck (OU) process. This enables precise control of the mean input (graded red triangle) and noise levels (inset) (See methods). **(D)** Raster plots illustrating a sharp transition from single spike to burst spikes with increasing dendritic inputs without noise. Noise leads to a more graded transition. Bursts are color coded in red. **(E)** Network configuration with distal dendrites of L5 PCs under control of inhibitory synaptic inputs from SOMs (blue circle). Bursts are activated by weak (pink, 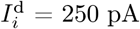) or strong (red, 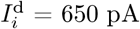) dendritic input with moderate noise levels (σ^d^ = 100 pA). The somatic input is the same for both dendritic inputs (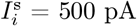, σ^s^ = 100 pA). The strength of the inhibitory connections w^SOM→dend^ is plastic (arrow) and modified according to our homeostatic plasticity rule (Eq. 1). The burst target rate (dashed line) was set to 1 Hz. **(F)** The burst rate after learning the inhibitory weights for different target burst rates. **(G)** Representative raster plots of the burst activity for weak and strong inputs, before and after learning. Each dot is a burst. **(H)** The distribution of the inter-burst intervals (IBI) before and after learning for weak and strong dendritic inputs.

### Controlling the burst activity of L5 pyramidal cells requires fine-tuning of the excitatory input and noise levels

The computational role of L5 PCs as bursting units depends on how dendritic and somatic inputs are translated into a spike and burst response. Due to the non-linear nature of the burst generation mechanism, we expected that the dynamic range of burst activity is limited when bursts are triggered by excitatory synaptic potentials on the apical dendrite alone^1^. In other words, the difference between no burst activity and saturated burst activity should be brought about by small differences in dendritic input. We checked this intuition in a population of model PCs by injecting current into both the soma and the dendrite. Spikes were generated by a noisy background input to the somatic compartment, with firing rates that mimic sensory driven activity^7^. The conversion of somatic spikes into bursts was driven by noisy excitatory current to the dendritic compartment, for which we systematically varied the mean and the noise level. We found that in the absence of noise on the dendritic input, an increase of the mean input to the dendrite leads to a rapid transition from an absence of bursts to a saturated level of burst activity (Fig 1C, light blue trace). The average population burst rate as a function of the dendritic input currents shows a step-like transition, at around 175 pA for our parameter settings. The majority of spikes appear as single spikes (singlets) below this threshold (Fig 1D, no noise condition). Above, all spikes are converted to bursts. The saturation level for bursting activity is determined by the amount of somatic input and potential refractory effects in the dendrite, which are mediated by a slow adaptation current that hyperpolarizes the dendrite after the sustained depolarization of a dendritic spike (see Methods). Hence, in the absence of noise in the dendrite, the non-linear dendritic threshold mechanism indeed limits the dynamic range of PCs as bursting units. Under this conditions, the low, consistent burst rate observed *in vivo*^2,3,7^ would require a fine-tuning of the input levels.

Noise can broaden the dynamic range of neural information transmission^26^. We therefore stimulated the dendrite with coloured noise with varying mean and variance (see Methods). Indeed, increasing dendritic noise changes the input-output relation between dendritic input and burst rate from all-or-none to a gradual transition (Fig 1C). Dendritic noise hence allows for a wider dynamic range for the possible burst rates and reduces the need to fine-tune input levels to achieve sparse bursting. A homeostatic control at low burst rates would therefore benefit from large fluctuations on the dendritic input currents. Large input fluctuations arise, e.g., in balanced networks, in which strong excitatory currents are on average cancelled by strong inhibitory currents^27,28^. Therefore, we next investigated if dendritic inhibition can mediate a control of burst activity and generate the fluctuations characteristic of a dendritic balanced state.

### Homeostatic inhibitory plasticity controls the burst rate of PCs

Neocortical SOM interneurons specifically target the distal tuft of pyramidal neurons^29^ and exert a profound influence on dendritic calcium activity and bursting^4,22^. To investigate if SOM interneurons can control the burst activity of pyramidal cells, we simulated a postsynaptic population of PCs receiving inhibitory input from SOM interneurons to the dendritic compartment (Fig 1E). Spikes in the SOM and PC population were generated by independent background noise, with firing rates that mimic sensory activity^30,31^.

We considered a burst timing-dependent homeostatic plasticity rule to regulate the strength of inhibitory synapses. In effect, synaptic efficacy is potentiated for near-coincident postsynaptic bursts and presynaptic spikes, while every presynaptic spike leads to synaptic depression. This burst-dependent rule is motivated by a previously proposed homeostatic plasticity rule designed to control postsynaptic firing rates^24^, but integrates post-synaptic bursts as salient plasticity-inducing events^9,13^. The learning rule can be summarised as

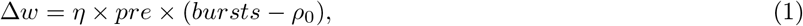

where *η* is the learning rate, *pre* is presynaptic activity, *bursts* is a postsynaptic trace reflecting recent burst activity, and *ρ*_0_ is a target rate for burst activity (see Methods for details). This rule is supported by experimental data insofar as inhibitory synapses from SOM interneurons onto CA1 pyramidal cells undergo potentiation when presynaptic activity is paired with postsynaptic bursts^32^.

We find that this learning rule robustly controls the burst rate of the postsynaptic neuron, both for high and for low excitatory input (Fig 1E, middle), by adjusting the synaptic weights of the inhibitory synapses onto the dendrite (Fig 1E, right). Homeostatic control is robust over a range of target rates covering both low burst rates and bursts rates near saturation (Fig 1F). The learning rule also controlled the temporal patterns of bursting. Before learning, burst activity is dense and sparse for strong and weak dendritic inputs, respectively. After learning, the PCs show similar burst raster plots (Fig 1G) and inter-burst interval (IBI) distributions (Fig 1H) for both initial conditions.

Because somatic burst activity may not be easy to sense for inhibitory synaptic connections on the apical dendrite, we wondered whether inhibitory synaptic plasticity in the dendrite could also be controlled by a postsynaptic signal local to the dendrites. Dendritic calcium spikes generate a long-lasting dendritic plateau potential (Fig S1A, red), which drives somatic bursting during BAC firing. Therefore, a thresholded version of the dendritic membrane potential provides a local estimate of the occurrence of a burst. Using this proxy for burst activity in the homeostatic inhibitory learning rule also leads to robust control of the burst rate (Fig S1), suggesting that homeostatic burst control could be achieved by a simple, biologically plausible mechanism. In the following, however, we will continue to use the burst-based implementation of the plasticity rule Eq. 1, because it allows for the interpretation of the target rate *ρ*_0_ as a burst rate.

Note that the dependence of the learning rule on presynaptic activity allows a stimulus-specific form of homeostatic control if the inhibitory interneurons differ in their stimulus tuning^24,33,34^. Because the interneurons form a homogenous population in the settings studied here, the presynaptic dependence in the associative term of the learning rule (pre × *bursts*) is not essential and can be dropped as, e.g. in^35^, without a qualitative change of the results (Fig S2).

### Simultaneous control of somatic and dendritic activity

Homeostatic inhibitory control of spiking activity has previously been demonstrated in simpler point neuron models without an explicit bursting mechanism^24,35^. Given the nonlinear interactions between soma and dendrite, we next wondered whether a simultaneous control of bursting activity and overall spiking activity could be achieved by somatic and dendritic inhibition. To this end, we extended the network model by a second class of inhibitory interneurons whose synapses target the somatic compartment of the PCs, akin to PV interneurons ^36^. Both PV and SOM populations are modeled as single compartment neurons, driven by external noisy inputs and provide inhibition to the PCs through current-based synapses (see Methods).

To control both somatic and dendritic activity, we have distinct rules for the two inhibitory connections. SOM→dendrite connections are subject to the plasticity rule in Eq. 1, while a different spike timing-dependent inhibitory plasticity rule^24^ in the PV→somatic connection controls the overall level of activity (Fig 2A). We find that both the spike rate and the burst rate reach their respective targets (Fig 2B,C left), but not necessarily in a monotonic fashion. For example, we observed a transient overshoot of burst activity when both firing rate and burst rate were initially too low (Fig 2B, top right). The underlying reason is that firing rate and burst rate are not independent. A decrease in somatic inhibition not only increases the firing rate, but also the burst rate. Hence, a homeostatic control of firing rate can transiently generate an overshoot in burst rate, which is only later corrected by an increase in dendritic inhibition (Fig 2B). The character of this transient effect is likely determined by the relative time scales of plasticity in the two synapse types, which in turn depend on the respective learning rates and the activity in the network.

**Figure 2.**
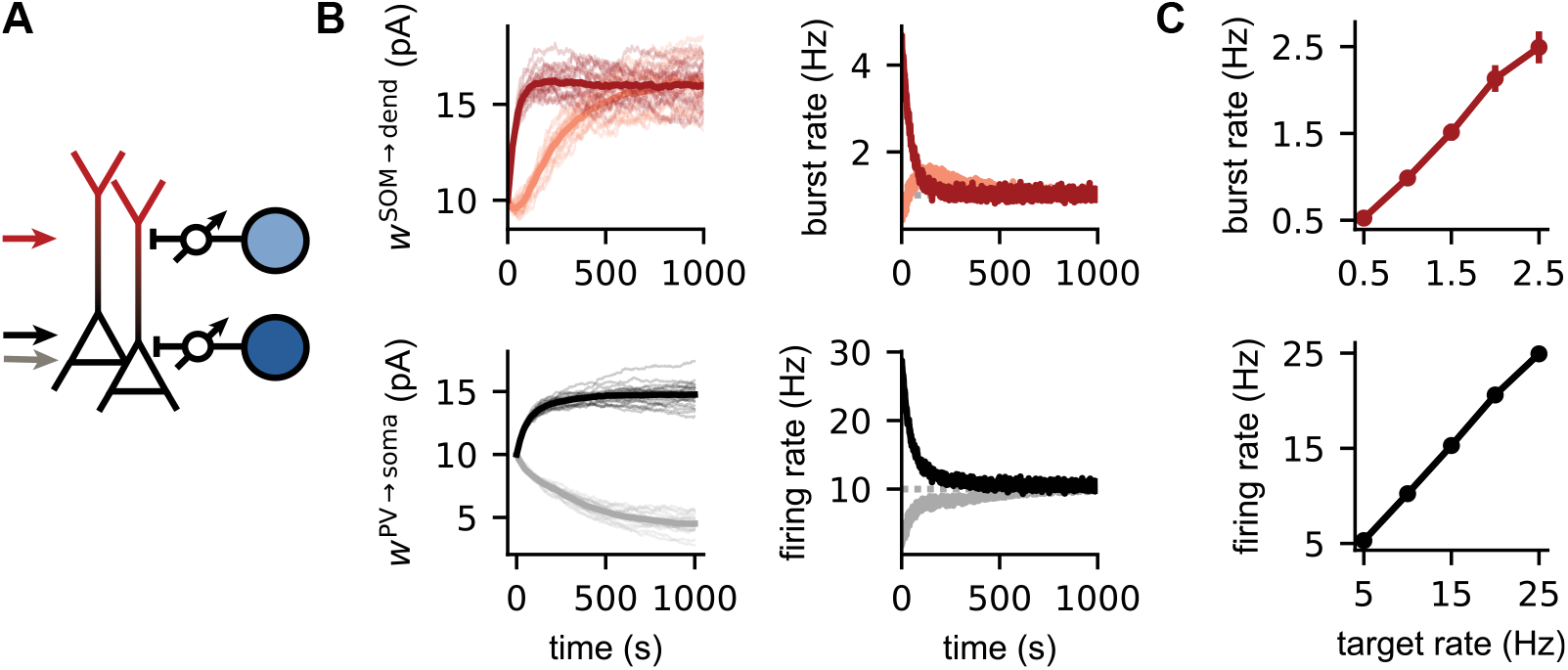
Simultaneous control of somatic and dendritic activity. **(A)** The somatic and dendritic activity of L5 PCs is under control of plastic inhibitory connections from PV (dark blue) and SOM (light blue) interneuron populations. The somatic compartment receives either weak (grey arrow, 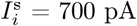) or strong (black arrow, 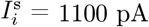) external inputs. The external input to the dendritic compartment is fixed (red arrow, 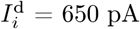). The variability of the noisy background input on the external inputs to the L5 PCs is moderate (σ^s^ = σ^d^ = 100 pA). **(B)** Evolution of the inhibitory weights, the firing and burst rates of L5 PCs during the learning process for strong (black/dark red) and weak (grey/light red) external somatic input. The target burst and firing rate are respectively 1 and 10 Hz. **(C)** The burst rate (top) and firing rate (bottom) after learning for different target rates for strong somatic 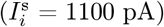 and dendritic input 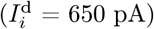. For all conditions, the target firing rate is 10 times larger than the target burst rate.

Homeostatic control is achieved over a range of target values for the firing rate and and the bursts rate (Fig 2C). Conflicts between the two learning rules only arise when the target for the firing rate is too low compared to the target for the burst rate (Fig S3). This is not surprising, because the firing rate introduces an upper limit for the burst rate. A firing rate of 10 Hz does not allow a burst rate higher than 5 Hz because bursts must by definition contain at least two spikes.

A simultaneous control of firing rate and burst rate can also be achieved in a recurrent microcircuit, in which the PV and SOM interneurons receive excitatory input exclusively from PCs (see next section, Fig 3). Thus, self-organised inhibition with local learning rules allows a precise control of somatic and dendritic activity in cortical microcircuits, by balancing somatic and dendritic excitation by suitable levels of inhibition.

**Figure 3.**
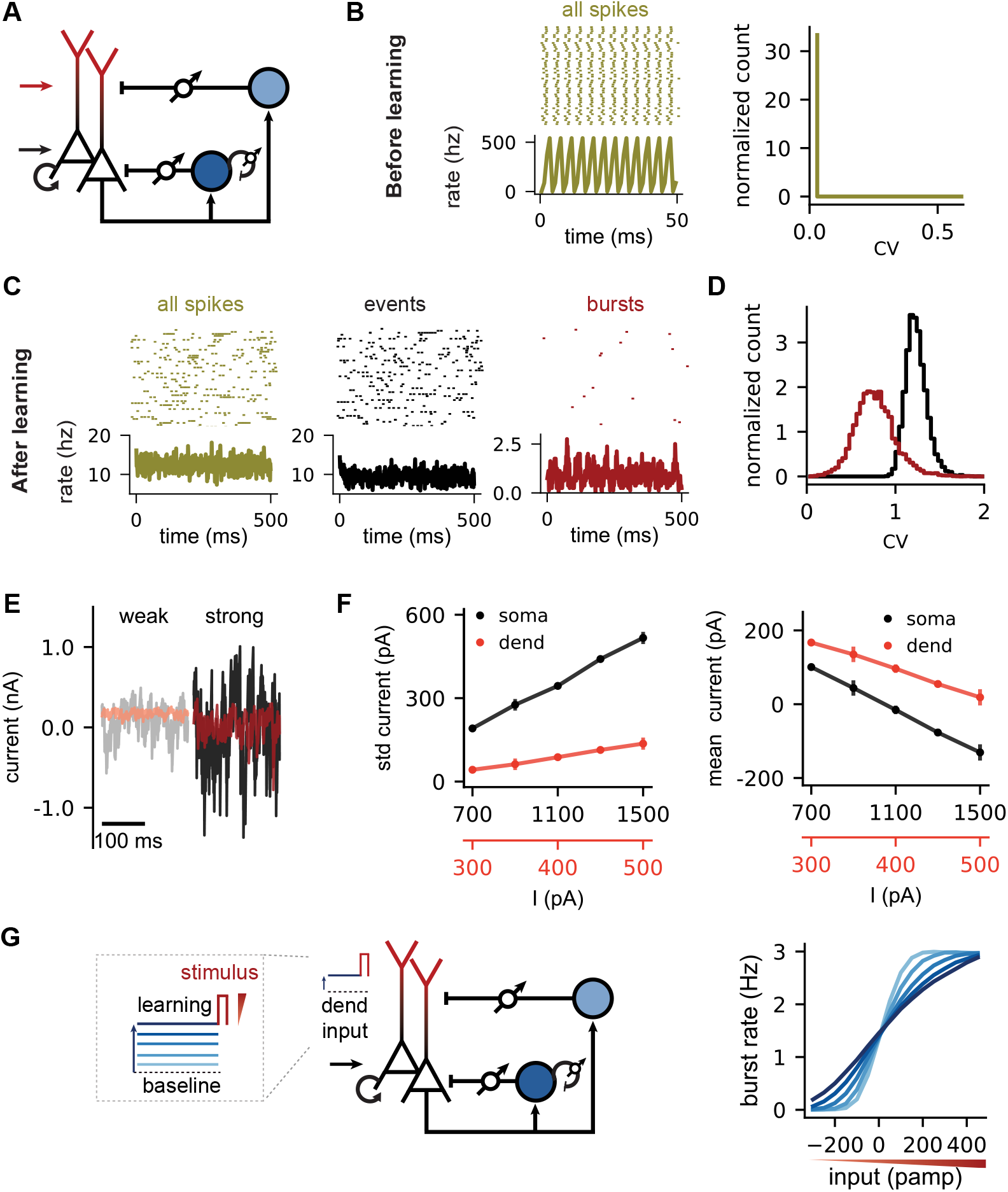
A doubly asynchronous irregular state for both spikes and bursts. **(A)** Dendritic and somatic activity of L5 PCs receive inhibition from respectively SOMs (light blue) and PVs (dark blue). In turn, L5 PCs drive SOMs and PVs. L5 PCs and PVs also receive excitatory and inhibitory recurrent connections respectively. All inhibitory connections are plastic (target firing rates = 10 Hz, target burst rate = 1 Hz). For all panels, the external input to soma, dendrites and PVs 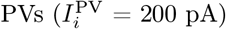 is modelled as constant input without noise (σ^s^ = σ^d^ = σ^PV^ = 0 pA). The SOMs do not receive external input. **(B)** Left: Spiking pattern of L5 PCs before learning of the inhibitory weights and time-varying population rate. Strong constant somatic 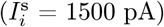 and dendritic 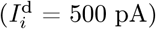 external input drives L5 PCs to fire synchronously at high rates without burst activity. Right: The firing pattern is regular as indicated by the distribution of the coefficient of variation (CV) of the inter-spike interval. **(C)** Raster plots and population rates after learning the inhibitory weights. Events (black) and bursts (red) are isolated from all spikes (dark yellow) to illustrate activity associated with somatic (events) and dendritic (bursts) regions (see Methods). **(D)** The firing and burst pattern is irregular as indicated by the distribution of the coefficient of variation (CV) of the inter-burst (red) and inter-event (black) intervals. **(E)** Net (exc + inh) input current to the soma (black traces) and dendrites (red traces). Weak (strong) external input leads to small (large) input fluctuations on the net input current. **(F)** Standard deviation (left) and the mean (right) net input currents for soma (black) and dendrites (red). **(G)** Left: Microcircuit and stimulation paradigm. During the learning phase, the inhibitory weights change until the somatic (10 Hz) and dendritic target (1.5 Hz) is reached for constant dendritic inputs ranging from weak to strong (blue 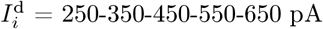). Right: After learning, the burst rate is computed during a transient input stimulus (red) and plotted as a function of the strength of the stimulus (red triangle). The somatic input (black arrow, 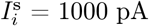) is the same for all dendritic input conditions.

### A doubly asynchronous irregular state for both spikes and bursts

Asynchronous irregular (AI) activity is a hallmark of recurrent networks in which excitation is balanced by inhibition, and can persist even in the absence of external noise sources^27,37^. Earlier work has shown that homeostatic inhibitory plasticity can establish such a fluctuation-driven AI state^24^. We therefore hypothesised that the combination of rate and burst homeostasis can self-organize a recurrent neural network into a doubly asynchronous irregular state for both spikes and bursts. We tested this hypothesis in a recurrent microcircuit in which all neurons in the circuit are driven by constant, noise-free external excitatory input (Fig 3A). We varied the strength of the input for the somatic and dendritic compartments of the PCs. In addition to plasticity of the inhibitory connections onto the PCs, we applied the homeostatic inhibitory plasticity rule to the inhibitory recurrent connections within the PV population to desynchronise the PV interneurons.

When we initialize the network with small inhibitory weights, the network initially synchronizes strongly at high firing rates (Fig 3B). At this point the absence of inhibition keeps the dendrites of the neurons in a persistently depolarised state, and an identification of bursts is pointless. Over the course of learning, inhibitory plasticity reduces both the firing rate and the burst rate to their respective targets, and the network develops asynchronous irregular activity patterns (Fig 3C). To assess the degree of irregularity of the activity without confounds from the presence of bursts, we studied the statistics not of individual spikes, but of events^17^. As events, we define individual spikes and the first spike within a burst (Fig 3C, black). Additional spikes within the burst are ignored. We find that both inter-event intervals and the inter-burst intervals are highly variable after learning (Fig 3D; mean CV of the inter-event interval distribution: 1.24; mean CV of inter-burst interval distribution: 0.77) indicating a doubly irregular state.

One hallmark of the fluctuation-driven, inhibition-dominated regime that underlies the AI state in balanced networks is that the mean input current within the population decreases with increasing external drive, while firing rates increase due to an increase in input variance. When we systematically varied the external drive to both the soma and the dendrites of L5 PCs, we find that this is also the case for both compartments (Fig 3E,F) in our network model. Note that the external inputs are noise-free, i.e., the variance in the inputs is generated intrinsically by the balance of excitation and inhibition, as for networks with simpler neuron models^27^. Functionally, this internally generated noise has the effect of smoothing out the input-output function of the network, such that transient inputs to the network can be represented in a graded rather than an all-or-none fashion (Fig 3G).

In summary, homeostatic inhibitory plasticity in somatic, dendritic and inter-interneuron connections can establish a doubly asynchronous irregular state in the network, in which both spikes and bursts occur irregularly, by means of internally generated noise.

### Inhibitory plasticity enables a multiplexed spike-burst code

While dendritic inputs to PCs are usually interpreted as “gain modulators” of PC responses^38–40^, spikes and bursts could also be used in a multiplexed ensemble code that allows to decode both the somatic and the dendritic input to a neuronal population^17^. According to this hypothesis, somatic input to the PCs is represented in the event rate of a population of PCs, while dendritic inputs are represented by the fraction of events that are bursts (burst fraction, BF; Fig 4A). Indeed, the event rate and burst fraction of a population of uncoupled PCs accurately tracked two time-varying inputs signals to their somata and dendrites (Fig 4B). However, this code is not robust to changes in input conditions. For the encoding of graded signals, the multiplexed code relies on noise in the input signals that smoothens out the neuronal input-output function and effectively decorrelates the responses of different neurons in the population. In other words, the population is artificially maintained in a fluctuation-driven regime by the addition of external noise. In line with this intuition, the decoding accuracy for both the somatic and dendritic input degrades when we add a constant baseline input to the dendrites (Fig 4C-E), shifting the neurons away from the fluctuation-driven and towards a mean-driven regime.

**Figure 4.**
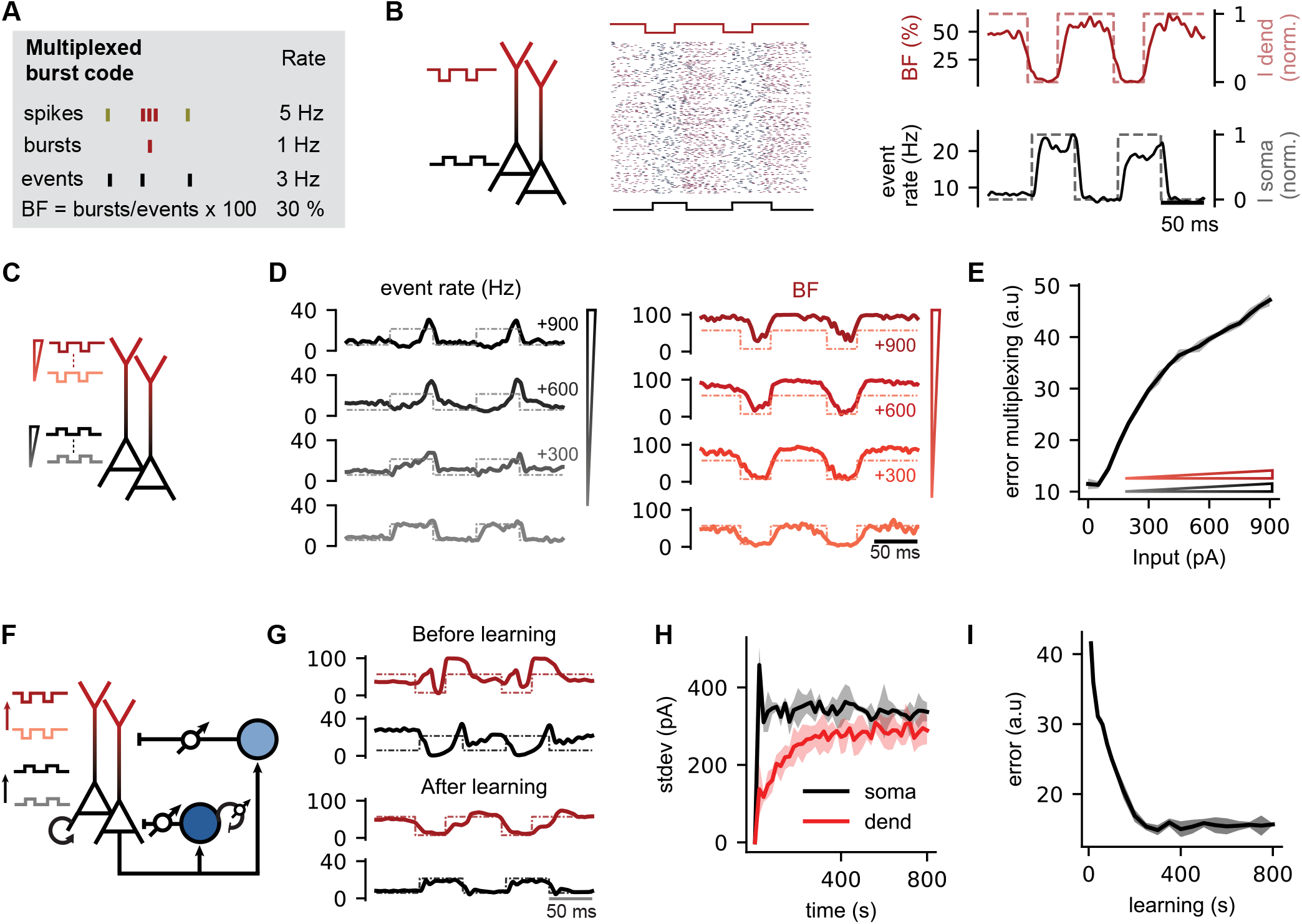
Inhibitory plasticity self-organises a multiplexed burst code. **(A)** Example illustrating a multiplexed burst code in L5 PCs in which somatic and dendritic inputs are represented in the event rate and the burst fraction, respectively^17^. Events are either a burst or single spike, while the burst fraction (BF) is the fraction of events that are bursts. **(B)** Alternating and opposite pulse inputs are delivered to the somatic and dendritic compartment 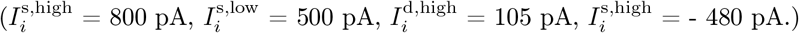 The event rate (black) and BF (red) respectively follow the somatic (dashed black) and dendritic input patterns (dashed red). **(C)** Stimulation paradigm with an increase in background excitation (triangle, red = dendrite, black = soma) on which pulse inputs are superimposed. **(D)** Compared to B, the dendritic and somatic background increases with 300, 600 and 900 pA, which deteriorates the multiplexed code. Dashed lines represent scaled versions of the alternating dendritic and somatic inputs (see Methods). **(E)** Quality of the multiplexed burst code as measured by the difference between the somatic (dendritic) input pulses and the event rate (BF) (see Methods). **(F)** Inhibitory plasticity restores the multiplexed burst code in a biological microcircuit without the need for fine-tuning the background input or noise levels. The microcircuit is similar to Fig 3, with constant external inputs, recurrent connections and plasticity on all inhibitory connections. Background excitation to both somatic and dendritic compartments was increased with 900 pA, the highest scaled input of Fig 4D, where event rate (BF) is not informative of somatic (dendritic) input pulses. **(G)** The event (black) and BF (red) before and after learning. Dashed lines represent scaled versions of the input. **(H)** Increasing inhibitory weights during the learning process increases the standard deviation of the net dendritic (red) and somatic (black) input currents. **(I)** Quantification of the quality of the multiplexed burst code while learning the inhibitory weights. For E,H and I, the solid line shows mean over 3 experiments with random network initialisation, while shaded region shows the 95 % confidence interval.

Given that the two forms of plasticity tend to establish a fluctuation-driven regime, we hypothesised that they could provide a basis for a self-organization of a multiplexed event-burst code. Indeed, we find that inhibitory plasticity in synapses from SOM neurons onto the dendrite can compensate the deleterious effects of a high baseline current to the dendrite (Fig 4F-I). SOM plasticity reduces the burst rate to a low, but consistent value (Fig 4G) and recovers a fluctuation-driven (Fig 4H), irregular asynchronous state that enables the multiplexed code (Fig 4G,I).

In summary, homeostatic plasticity of inhibitory synapses can put neural circuits with complex PC dynamics and two different interneuron classes into a fluctuation-driven regime, which not only establishes a doubly balanced state, but also stabilizes a multiplexed neural code for bottom-up and top-down signals.

## Discussion

The question of how pyramidal neurons integrate bottom-up and top-down information streams has received keen interest over the past decades. Here, we addressed the question how a network can self-organize into a dynamical state in which this integration is likely to be most effective. We have shown i) that a simple form of inhibitory plasticity can homeostatically control the burst rate, ii) that it can be readily combined with a homeostasic control of firing rate, iii) that this form of homeostasis can establish a doubly irregular network state for spikes and bursts, and iv) that this state indeed improves the ability of bursts to convey information in a multiplexed neural code^17^.

### Functional benefits of homeostatic burst control

Given the broad range of potential dendritic computations ^41^, a homeostatic control of dendritic and/or burst activity could serve a variety of functions. Sparse bursting maximizes information in multiplexed neural codes. Such multiplexed codes could in turn allow a bi-directional propagation of signals in cortical hierarchies^17^, e.g., a backpropagation of internal predictions^11,12,42^ or error signals^13,18^ to lower layers in the hierarchy.

In line with the notion that bursts could represent error signals, they are very effective drivers of synaptic plasticity^9^, suggesting that learning can be regulated or at least influenced by inputs to the upper cortical layers. Notably, error-driven learning is substantially more effective when the error signals are graded rather than all-or-none. Therefore, the suggested homeostatic control of burst activity with the accompanying response linearization may be beneficial to create suitable conditions for graded learning signals ^13^. Finally, if plasticity is primarily triggered by bursts, a homeostatic control of burst activity could be interpreted as a form of “meta-plasticity” that controls how often plasticity is triggered in a given neuron or circuit. A similar argument can be made for the idea of coincidence detection by somato-dendritic integration^10^. Homeostatic inhibitory plasticity in the dendrite could serve to set a (potentially soft) threshold above which dendritic input is deemed sufficiently high to trigger the coincidence detection machinery.

A different argument for homeostatic inhibitory plasticity is the establishment of a balanced state^43,44^, in which excitation and inhibition cancel out on average^27,37^. While the energy expenditure of balanced states is a frequent target of mockery but see^45^, the underlying idea of inhibitory negative feedback loops has the advantage of smoothing out threshold-like processes and thereby broadening the dynamic range for information transmission^46^ (Fig 1). In line with this idea of response linearization, responses to dendritic stimulation are more graded in vivo than *in vitro* ^22^.

### Specificity of homeostatic control

A frequent question for excitation-inhibition (E/I) balance is that of its spatiotemporal precision, i.e., the question along which dimensions excitation and inhibition are correlated and how tight this correlation is^43^. Originally suggested as a balance on the network level and merely present on average across time and neurons^27^, the E/I balance can also be specific in time^28,47^, in stimulus space^48,49^, across neurons^50^ or across neuronal compartments^51,52^. Each of these dimensions of specificity has its correspondence in potential inhibitory learning rules that could establish the respective form of E/I balance. Specificity across neurons requires a dependence of inhibitory plasticity on postsynaptic signals ^24,35^. Specificity in time and stimulus requires a dependence on presynaptic activity^44,53^. A specificity across compartments — as studied here — requires a dependence on compartment-specific signals. In our simulations, we used bursts as a proxy for dendritic activity in L5 pyramidal neurons, but membrane potential (Fig S1) or currents in the dendrite or local chemical signals could be equally suitable. We included a dependence on presynaptic activity in the dendritic learning rule to leave open the possibility of both a compartment- and input-specific E/I balance in further studies. However, in the situations studied here, the presynaptic interneuron populations are homogeneous, so the presynaptic dependence does not have an impact on the results (Fig 1 versus Fig S2).

### Experimental support & interaction of homeostatic mechanisms

A key prediction of the model is that inhibitory synapses from SOM interneurons onto PCs should undergo potentiation when the postsynaptic cell bursts too often. This is supported by slice experiments in the hippocampus^32^, which showed that a theta burst stimulation protocol – presynaptic activity paired with regularly occurring postsynaptic bursts – induces long-term potentiation in SOM→PC synapses. Notably, the same protocol induces long-term depression in PV→PC synapses. Different interneurons hence display different rules of synaptic plasticity – also on their excitatory input synapses^54^–, which rely on distinct molecular mechanisms^32,55^.

In our simulations, the effects of different forms of homeostatic plasticity are not necessarily independent. Homeostatic control of the overall firing rate also influenced the rate of bursts (Fig 2), because bursts generated via BAC firing^4^ are triggered by somatic spikes and hence depend on the firing rate. Such interactions arise when the sensors (firing rate/burst rate) or the effects (SOM→PC/PV→PC synapses; somatic/dendritic membrane potential) of the homeostatic control laws are correlated, and can generate non-monotonic homeostatic dynamics (Fig 2B) (similar behavior was seen, e.g., by O’Leary et al.^56^). This creates potential challenges for an experimental investigation of dendrite-specific forms of homeostasis.

The basic prediction of our model is that an over-activation or suppression of dendritic activity should result in specific compensatory changes in inhibitory synapses onto the dendrite. However, it may not be trivial to manipulate dendritic activity without manipulating other aspects of network activity. Depending on the experimental manipulation (e.g., tissue-wide application of TTX^57^ vs. application of gabazine or baclofen to the superficial layers^22^) and the observed quantities (burst rate, dendritic calcium signals or morphological features of inhibitory synapses on the dendrite), the observations could differ substantially, if other homeostatic mechanisms occur in parallel. Moreover, it is conceivable that different forms of homeostasis interact to decorrelate their effects. For example, a neuron could react to high dendritic activity by redistributing inhibition from the soma to the dendrite, in order to selectively reduce bursts without affecting overall firing rate. A stimulation protocol based on postsynaptic bursts would then simultaneously potentiate dendritic inhibition and depress somatic inhibition. The observed opposing forms of plasticity in SOM and PV synapses for the same stimulation protocol^32,55^ could therefore be interpreted as a decorrelation of the effects of these two synapse types on firing rate and burst rate.

### What’s wrong or missing in the circuit

The primary focus here was on the self-organisation of a dynamical network state in which bursts occur rarely. Like all models, we navigated a trade-off between model simplicity, clarity of result and biological accuracy, and the circuit we studied is clearly simplified compared to cortical circuits. For simplicity, we used the same low connection probability among all neuron classes, although the connection probability of interneurons is substantially higher than that of excitatory neurons^58,59^. We expect that the key results carry over denser connectivity, despite potentially higher input correlations^28^.

Several cortical interneuron classes were ignored, including interneuron types that also inhibit the distal dendrite^20,60^. Those interneurons could provide the suggested homeostatical control of dendritic activity equally well as SOM interneurons. Whether two distinct interneuron classes are actually required for a compartment-specific form of feedback inhibition or whether this could be mediated by a single cell class with heterogeneous properties was investigated elsewhere^52^. We also ignored the well-documented connections between SOM and PV neurons^19,61^. In the presence of stimuli, these connections could mediate a redistribution of inhibition across the two compartments^62^, but in the steady-state conditions we studied here, they would likely not change the results qualitatively. Additional interneurons that mediate – e.g., a dynamic disinhibition of the dendritic compartment^62–66^ – would also become relevant players in the presence of time-varying inputs.

### Outlook

Natural extensions of this work would be the addition of time-varying or stimulus-dependent input, combined with a stimulus tuning of the various cell classes, to study simultaneously the effects of stimulus-specific ^24,33,34^ and compartment-specific^52^ E/I balance. To do so, however, we would have to specify a stimulus selectivity for all neuron classes in the network ^33,34^ and the resulting rich combinatorics of conditions is beyond the scope of this work.

## Methods

### Network model

We gradually increase the complexity of the network from an uncoupled population of PCs to a feedforward network with inhibition and, finally, a recurrent network with two interneuron classes, representing dendrite-targeting SOM interneurons and soma-targeting parvalbumin-positive (PV) interneurons. All neurons are randomly connected. Parameters are provided in Tables 1, 2 & 3.

**Table 1.** Parameter values for the two-compartmental L5 PC model. Soma and dend indicate the somatic and dendritic compartment respectively and *f*(*x*) the sigmoid function. Values are from^17^.

| soma | | | dendrite | | | $f(x)$ | | |
| --- | --- | --- | --- | --- | --- | --- | --- | --- |
| $\tau^s$ | 16 | ms | $\tau^d$ | 7 | ms | $E_d$ | -38 | mV |
| $C^s$ | 370 | pF | $C^d$ | 170 | pF | $D_m$ | 6 | mV |
| $g^s$ | 1300 | pA | $g^d$ | 1200 | pA | | | |
| $b_w^s$ | -200 | pA | $c^d$ | 2600 | pA | | | |
| $\tau_w^s$ | 100 | ms | $\tau_w^d$ | 2600 | pA | | | |
| $E_L$ | -70 | mV | $a_w^d$ | -13 | nS | | | |
| | | | $E_L$ | -70 | mV | | | |

**Table 2.** Parameters of the PV and SOM interneuron models.

| PV |  |  | SOM |  |  |
| --- | --- | --- | --- | --- | --- |
| $\tau^{\text{PV}}$ | 10 | ms | $\tau^{\text{SOM}}$ | 20 | ms |
| $C^{\text{PV}}$ | 100 | pF | $C^{\text{SOM}}$ | 100 | pf |
| | | | $b_w^{\text{SOM}}$ | -150 | pA |
| | | | $\tau_w^{\text{SOM}}$ | 100 | ms |

**Table 3.** Number of neurons in each populations for the different figures.

| FIGURE | 1 | 2 | 3 | 4 |
| --- | --- | --- | --- | --- |
| $N^{\text{PC}}$ | 1600 | 1600 | 8000 | 8000 |
| $N^{\text{SOM}}$ | 400 | 400 | 2000 | 2000 |
| $N^{\text{PV}}$ | - | 400 | 2000 | 2000 |

#### L5 PCs

L5 PCs are simulated as a two-compartmental model akin to the model described by Naud et al.^17^. The two compartments represent the soma and distal dendrites, and their interaction captures dendrite-depending bursting. The membrane potential *V^s^* of the somatic compartment follows generalised leaky integrate-and-fire dynamics with spike-triggered adaptation. The subthreshold dynamics of the *i*^th^ pyramidal neuron is described by

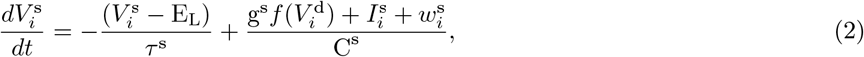

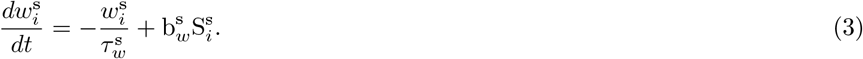

The dynamics of the somatic membrane potential 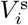 are governed by a leak term that drives an exponential decay to a resting membrane potential E_L_ with membrane time constant *τ*^s^. When the somatic membrane potential reaches a threshold V_T_, it is reset to the reversal potential after an absolute refractory period of 3 ms and a spike is added to the spike train 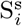. The soma is subject to spike-triggered adaptation. Each somatic spike increases an adaptation current *w^s^* by an amount 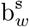. Between spikes, the adaptation current *w^s^* decays exponentially with time constant 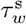. The soma receives external inputs 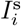 and a current 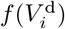 from the apical dendrite that depends nonlinearly on the dendritic membrane potential 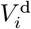. The parameter g^s^ controls the coupling strength of the dendrite to the soma. The impact of all these currents on the somatic membrane potential is scaled by the somatic membrane capacitance *C*^s^.

The dendritic compartment is modeled by the following dynamics:

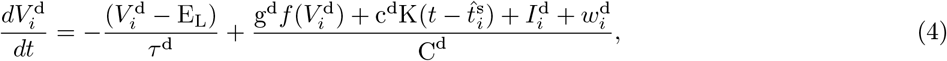

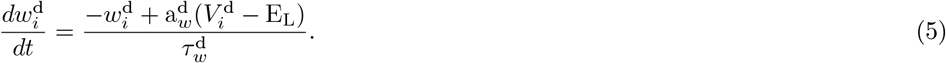

The dendritic membrane potential 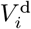 decays expontentially to the resting membrane potential E_L_, with a time constant *τ*^d^. Dendritic calcium events are modeled as a nonlinear current 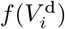, which increases steeply when the dendritic membrane potential approaches a given threshold E_d_:

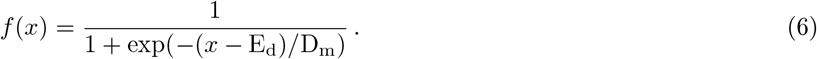

The steepness of this threshold is controlled by the parameter D_m_. The coupling from the somatic to the dendritic compartment by backpropagating action potentials (BAPs) is modelled by a pulse-shaped current in the dendrite. The strength of this current pulse is controlled by the parameter c^d^ and its shape by a kernel *K*. *K* is a rectangular kernel of amplitude one, which lasts 2 ms and is delayed by 0.5 ms relative to the time 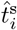 at which the somatic spike occurred. The dendrite is subject to subthreshold adaptation, which terminates dendritic calcium events unless external currents do so. The dynamics of the dendritic adaptation variable are defined by a strength 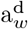 and time constant 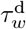. Again, all currents to the dendritic compartment are scaled by the dendritic membrane capacitance C^d^.

For Fig 3 we removed the spike-triggered somatic adaptation *w^s^* (Eq. (3)) to allow for short interspike intervals (ISI) between somatic spikes. This was necessary to achieve a CV of the interspike interval close to 1, because adaptation typically causes more regular activity by reducing the occurrence of short interspike intervals^67^. However, removing the somatic adaptation variable 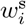 from Eq. (2) prolongs dendritic calcium spikes and bursts. To compensate this effect, the value of the dendritic adaptation variable 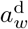 was increased from −13 pA to −28 pA to shorten bursts.

#### PV and SOM interneurons

The dynamics of the two interneuron populations are modelled by integrate-and-fire neurons. The subthreshold voltage dynamics 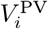 of the *i^th^* PV neuron is described by

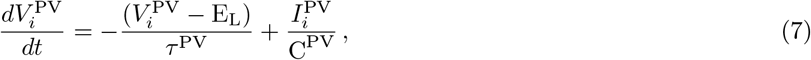

with membrane time constant *τ*^PV^ and capacitance C^PV^.

In contrast to PVs, SOMs exhibit firing rate adaptation 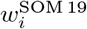, and are therefore described by an adaptive integrate-and-fire model,

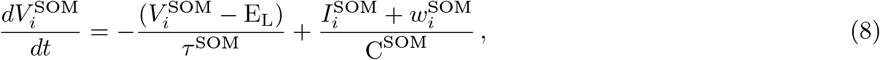

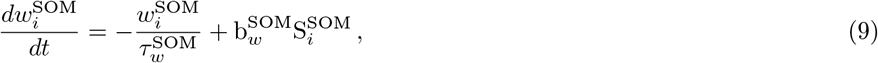

where *w*^SOM^ increases by 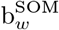 in case of a spike 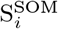 and decays otherwise at a rate defined by 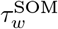. While PVs have a membrane time constant *τ*^PV^ of 10 ms, SOMs are modelled with a longer time constant 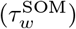 of 20 ms to be consistent with *in vivo* measurements^68^. The parameters *τ*^SOM^ and C^SOM^ are the membrane time constant and capacitance of SOMs, respectively.

### Connectivity

Specific network configurations were used for the different figures. The number of neurons in the network for the main figures of the manuscript are summarized in Table 3. Figs S1&S2 have the same network configuration as Fig 1, while Fig S3 and Fig S4 have the same network configuration as Fig 2 and Fig 3, respectively. All neurons are fully connected in Figs 1&2 (connection probability = 1) and with sparse random connectivity in Figs 3&4 (connection probability = 0.02). The network diagram depicted in each figure specifies the synaptic connections between the different neuron populations, with arrows indicating excitatory connections and straight arrow heads indicating inhibitory connections. All cells have the same number of incoming connections (homogeneous network), while autapses were excluded from the recurrent inhibitory PV_*i*_ → PV_*j*_ connections. The excitatory connections in Figs 3&4 were not plastic and the synaptic weights *w_ij_* fixed for 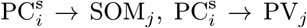 and 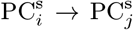. The strengths of the synaptic weights 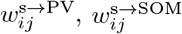 and 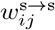 are 25 pA for Fig 3, while for Fig 4 the values are 15, 4 and 13 pA respectively (see also section on Inputs). All inhibitory connections were plastic and evolved according to the inhibitory plasticity rules. The inhibitory weights for Figs 1&2 were initialised at 10 pA while for Figs 3&4 inhibitory weights were initialised at 0.1 pA.

### Inhibitory plasticity

Spiking activity of L5 PCs is regulated by an inhibitory plasticity rule described in ^24^. Inhibitory synapses are strengthened by coincident pre- and postsynaptic activity within a symmetric coincidence time window of width *τ*_STDP_ (= 20 ms). Additionally, every presynaptic spike leads to a reduction of synaptic efficacy. In order to calculate the changes to each *W_ij_*, a synaptic trace x¿ is assigned to each neuron and *x_i_* increases with each spike *x_i_* → *x_i_* + 1. Otherwise it decays following

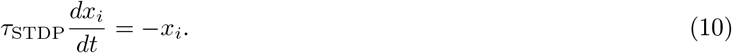

The synaptic weight *W_ij_* from neuron *j* to neuron *i* is updated for every pre- or postsynaptic event such that

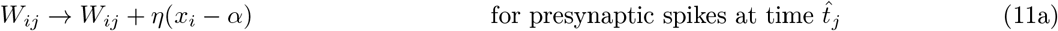

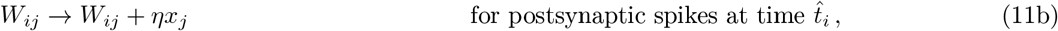

where *η* is the learning rate, *α* = 2 × *ρ*_0_ × *τ*_STDP_ the depression factor, and *ρ*_0_ the target firing rate.

The burst activity of L5 PCs is controlled by an analogous inhibitory plasticity rule as described above. Again, presynaptic activity *j* is captured by a synaptic trace that increases with each spike. The postsynaptic activity corresponds to burst activity and requires a different implementation. We explored two different strategies, an algorithmic (Figs 1, 2, 4, S2 and S3) and a voltage-based strategy (Figs 3 and S1). The algorithmic rule increases the postsynaptic trace *i* for each somatic burst (see below for our classification criteria for bursts), and decays otherwise with the time constant *τ*_STDP_ (= 20 ms), following Eq. (10). The synaptic weight *W_ij_* from neuron *j* to neuron *i* is updated for every pre- or postsynaptic event as in Eqs. (11). Similar to the inhibitory rule described in^24^, the algorithmic rule mathematically relates a target burst rate *ρ*_0_ with the target *α* by *α* = 2 × *ρ*_0_ × *τ*_STDP_. Note that we used different targets for the two inhibitory plasticity rules. The algorithmic implementation permits updates of the inhibitory weights as an explicit function of the burst rate, but requires somewhat “non-local” somatic information to update dendritic inhibitory weights. We therefore implemented an alternative rule to demonstrate that the burst rate can also be controlled using local dendritic signals. In this implementation, the post-synaptic activity is represented by dendritic calcium spikes. We approximate dendritic calcium spikes by thresholding the voltage of the dendritic compartment using a sigmoid function (Eq.(6)) with a sharp threshold at −20 mV (E_d_ = −20 mV and D_m_ = 0.01 mV). The relationship between the target *α* and the burst rate is determined empirically by plotting the burst rate as a function of increasing target values (see Fig S1). *W_ij_* is updated every time there is a pre- or postsynaptic spike.

The spike-time dependent plasticity rule controlling burst activity can be simplified further to a spike timing-independent model (Fig S2). In this rule, the changes to *W_ij_* do not require coincident pre- and postsynaptic activity and update such that

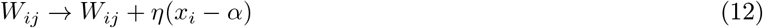

where *η* is the learning rate and *x_i_* is a trace representing postsynaptic burst activity. Similar as described above, an algorithmic and a voltage-based strategy can increase the postsynaptic trace *x_i_*, which decays otherwise with the time constant *τ*_STDP_, following Eq.(10). The algorithmic rule relates the target rate *ρ*_0_ for bursts to the depression parameter *α* by *α* = *ρ*_0_ × *τ*_STDP_ and was used in Fig S2). The synaptic weights are updated with a fixed regular time interval of 50 ms.

The learning rate *η*^SOM→d^ is 0.1 for Fig 1, S1 and S2. For Fig 2 and S3, the learning rates *η*^SOM→d^ and *η*^PV^→^s^ are 0.1 and 0.01, respectively. For Fig 3 and S4, the learning rates *η*^SOM^→^d^, *η*^PV^→^s^ and *η*^PV^→^PV^ are 1, 2.5 and 0.075, respectively, and 1, 0.1 and 0.05 for Fig 4.

### Inputs

The input to the neurons is characterised by external constant input *I*^ext^, noisy background input 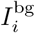 and synaptic input 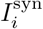:

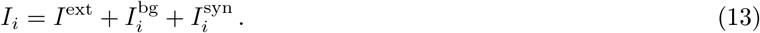

The noisy background input 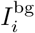 is modeled as an Ornstein-Uhlenbeck process with mean *μ*, variance *σ*^2^ and correlation time *τ*^OU^

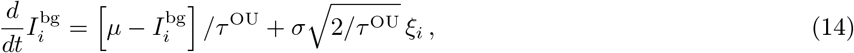

where *ξ_i_* is Gaussian white noise with 〈*ξ_i_*〉 = 0 and 〈*ξ_i_*〉(*t*)*ξ*_*i*_^(^*t*′)〉 = *δ*(*t* – *t*′). For all simulations *μ* and *τ*^OU^ were 0 pA and 2 ms respectively. The parameters of the external input *I*^ext^ and the standard deviation of the background input (σ) to L5 PCs are specified in the caption of each figure. The external and background inputs for the inhibitory SOM and PV populations in Fig 1&2 was chosen so that the firing rates were 10 Hz (Fig 1: *I*^ext,SOM^= 90 pA, *σ*^SOM^ = 400 pA; Fig 2: *I*^ext,SOM^ = 90 pA, *I*^ext,PV^ = −45 pA, *σ*^SOM^ = *σ*^PV^ = 400 pA).

The total synaptic input 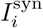 is the sum over all synaptic input currents triggered by all presynaptic neurons where the *f*-th presynaptic spike time of neuron *j* is labeled 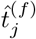:

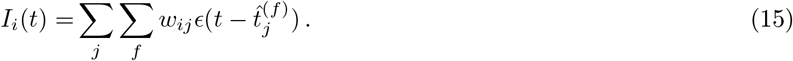

The time course of the synaptic input is modelled as an instantaneous jump followed by an exponential decay with a time constant of *τ* = 5 ms for excitatory synapses and *τ* = 10 ms for inhibitory synapses,

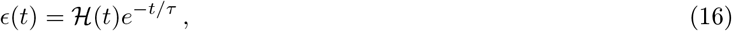

where 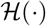 is the Heaviside step function and *w_ij_* the synaptic weight.

### Data Analysis

#### Neurometric parameters

Bursts are defined as a set of spikes where the interspike interval (ISI) is smaller than 16 ms. For Fig 3, in addition to the 16 ms ISI requirement, the presence of a dendritic calcium spike was verified to identify bursts. This was necessary because the absence of adaptation in the somatic compartment (Eq 3) can lead to ISIs below 16 ms, even without a dendritic calcium spike. See section on the inhibitory plasticity rules (voltage based strategy) for the identification of a dendritic spike. Events are all isolated spikes and the first spike of a burst. Burst and event rate are calculated by summing the bursts and events across the population, respectively. Burst probability is calculated as the ratio of the burst rate over the event rate. The time-dependent rates are smoothed for display, unless otherwise stated, by convolving the neurometric parameters burst and event rate with a rectangular window. The window-length is 2.5 seconds for Fig 1, Fig 2, Fig S1, Fig S2 and 10 ms for Fig 4. The rates for Fig 3&S4 are not smoothened to be able to evaluate the population rate at a temporal resolution in the ms range. So population rates are computed by counting the the total number of spikes of the whole population per 1 ms time-bins, and normalize the counts by the number of L5 PC neurons (8000) and the bin size (1 ms).

#### Coefficient of variation

To characterize the global state of the network we monitored the interspike intervals of individual spike trains. A hallmark of cortical activity is irregular asynchronous network activity and has a coefficient of variation of interspike intervals (ISI CVs) near 1^26,37^. ISI CV values close to zero indicate regular spiking patterns, values near 1 indicate irregular spiking. However burst activity confounds the interpretation of CVs since bursts can increase the CV independent of spiking regularity. To interpret the CV independent of burst activity we quantifies the regularity of events (IEI CVs). The regularity of bursts is quantified by computing the coefficient of variation of the inter burst intervals (IBI CVs).

#### Multiplexing error

In a multiplexed burst code spikes are separated in bursts and events to recover the input streams that arrive at the somatic and dendritic compartments of L5 PCs^17^ (see Fig 4A). The encoding quality of the dendritic and somatic input signals in a multiplexed burst code (see Fig 4E,I) is measured by comparing the shape of dendritic input 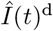 with the shape of burst fraction (BF(t)) and the shape of somatic input 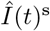 with the shape of event rate (ER(t)). This requires rescaling of the somatic and dendritic input currents (in units of pA) to match the scale of event rate (in Hz) and BF (unitless). Specifically, the minimum and maximum value of somatic and dendritic inputs were rescaled to match with the minimum and maximum values of event rate and BF, and computed as follows:

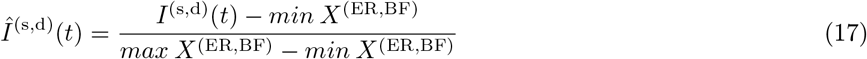

where, 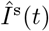 indicates the rescaled somatic input, 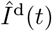 the rescaled dendritic input and X either the event rate (ER) or the burst fraction (BF). Examples of rescaled inputs are shown as dashed input pulses (grey for somatic input, and light red for dendritic inputs) in Fig 4D,G.

The encoding quality of the dendritic and somatic input is quantified by summing the absolute difference over all time points t between BF(t) and the rescaled dendritic input on the one hand and the difference between ER(t) and the rescaled somatic input on the other hand. Averaging both values quantifies the ‘error multiplexing’

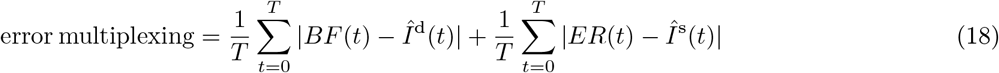

with 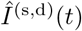 the rescaled version of the somatic or dendritic input current and T the length of the stimulation interval.

### Simulation details

All simulations were performed using the Brian simulator version 2.2.2.1^69^. Differential equations were numerically integrated using the Euler integration method with a time step of 0.1 ms. Source code to illustrate control of somatic and dendritic activity, i.e. Fig 1 and 2, will be available after publication.

## Acknowledgments

We thank Joram Keijser for discussions and feedback on the project, and Owen Mackwood for helpful discussions on network implementation and simulations. They, along with Loreen Hertäg, Laura Naumann and Felix Lundt also provided careful proof-reading of the manuscript. This project was funded by the Einstein Foundation Berlin via an Einstein Project to HS and Matthew Larkum (grant 1-4000015-01-EF) and by the German Federal Ministry Education and Research via a Bernstein Award to HS (grant 01GQ1201).

## Contributions

F.V., R.N. and H.S conceived the model and contributed to the interpretation of the results. F.V. performed the simulations. H.S. supervised the project, and acquired the funding. F.V. and H.S. wrote the manuscript.

## Supplementary figures

**Figure S1.**
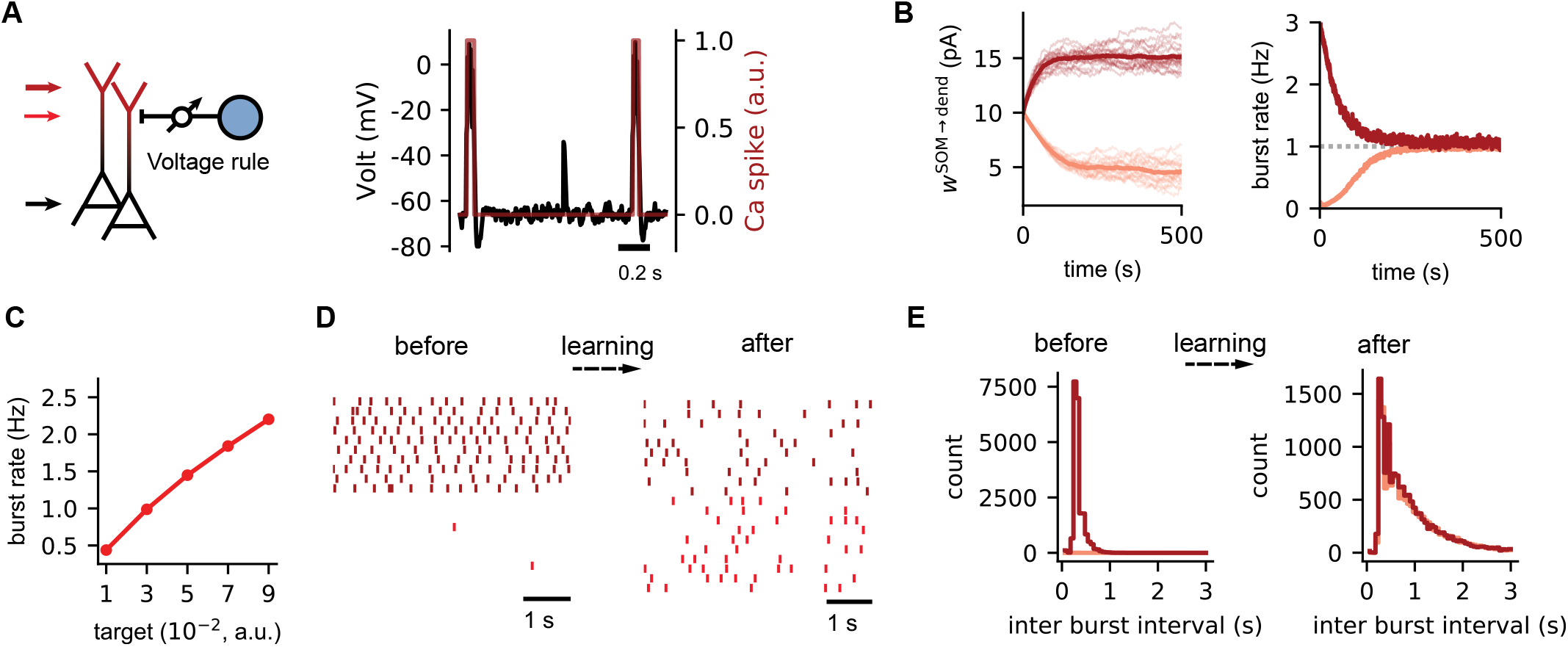
Control of the burst rate by a voltage based homeostatic inhibitory plasticity rule. **(A)** Network configuration with distal dendrites of L5 PCs under control of inhibitory synaptic inputs from SOMs (blue circle). The inhibitory connections are plastic (arrow) and modified according to a homeostatic plasticity rule where post-synaptic activity is modelled by a filtered version of the dendritic voltage (right, red trace)(methods). **(B)** Bursts are activated by weak (light red, 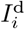 250 pA) or strong (dark red, 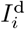 650 pA) dendritic input with moderate noise levels (σ^d^ = 100 pA). The somatic input is the same for both dendritic inputs (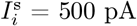, σ^s^ = 100 pA). The target value was determined empirically (see C) so that the burst rate was 1 Hz (dashed line). **(C)** The burst rate after learning the inhibitory weights for different target values. **(D)** Representative raster plots of the burst activity for weak (light red) and strong (dark red) dendritic inputs, before and after learning. Each dot represents a burst. **(E)** The distribution of the inter-burst intervals (IBI) before and after learning for weak (light red) and strong (dark red) dendritic inputs.

**Figure S2.**
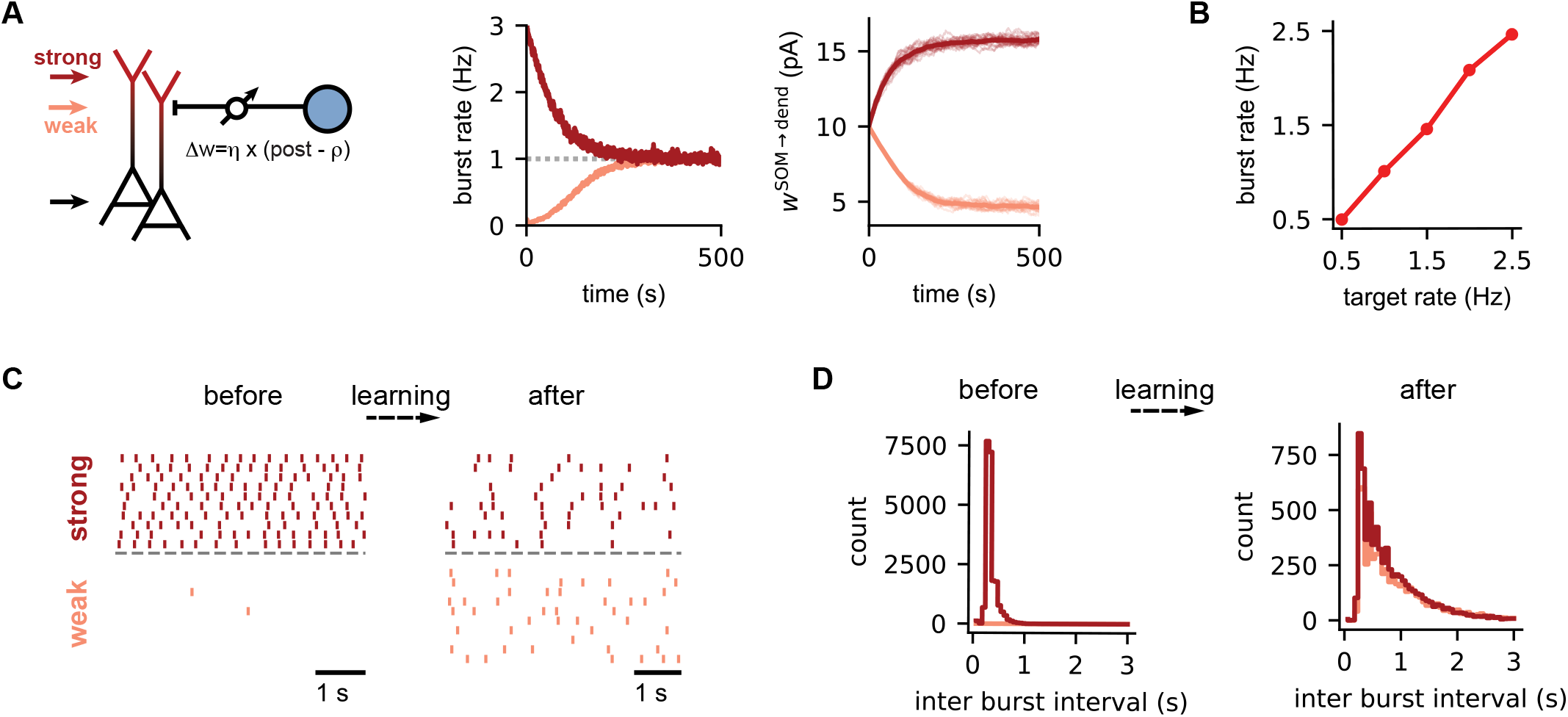
Control of the burst rate by spike-timing-independent homeostatic inhibitory plasticity. **(A)** Network configuration with distal dendrites of L5 PCs under control of inhibitory synaptic inputs from SOMs (blue circle). Bursts are activated by weak (light red, 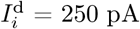) or strong (dark red, 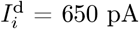) dendritic input with moderate noise levels (σ^d^ = 100 pA). The somatic input is the same for both dendritic inputs (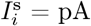, σ^s^ = 100 pA). The strength of the inhibitory connections W^SOM→dend^ are plastic (arrow) and modified according to a homeostatic plasticity rule dependent on dendritic post-synaptic activity (methods). The burst target rate (dashed line) was set to 1 Hz. **(B)** The burst rate after learning the inhibitory weights for different target burst rates. **(C)** Representative raster plots of the burst activity for weak (light red) and strong (dark red) dendritic inputs, before and after learning. Each dot represents a burst. **(D)** The distribution of the inter-burst intervals (IBI) before and after learning for weak (light red) and strong (dark red) dendritic inputs.

**Figure S3.**
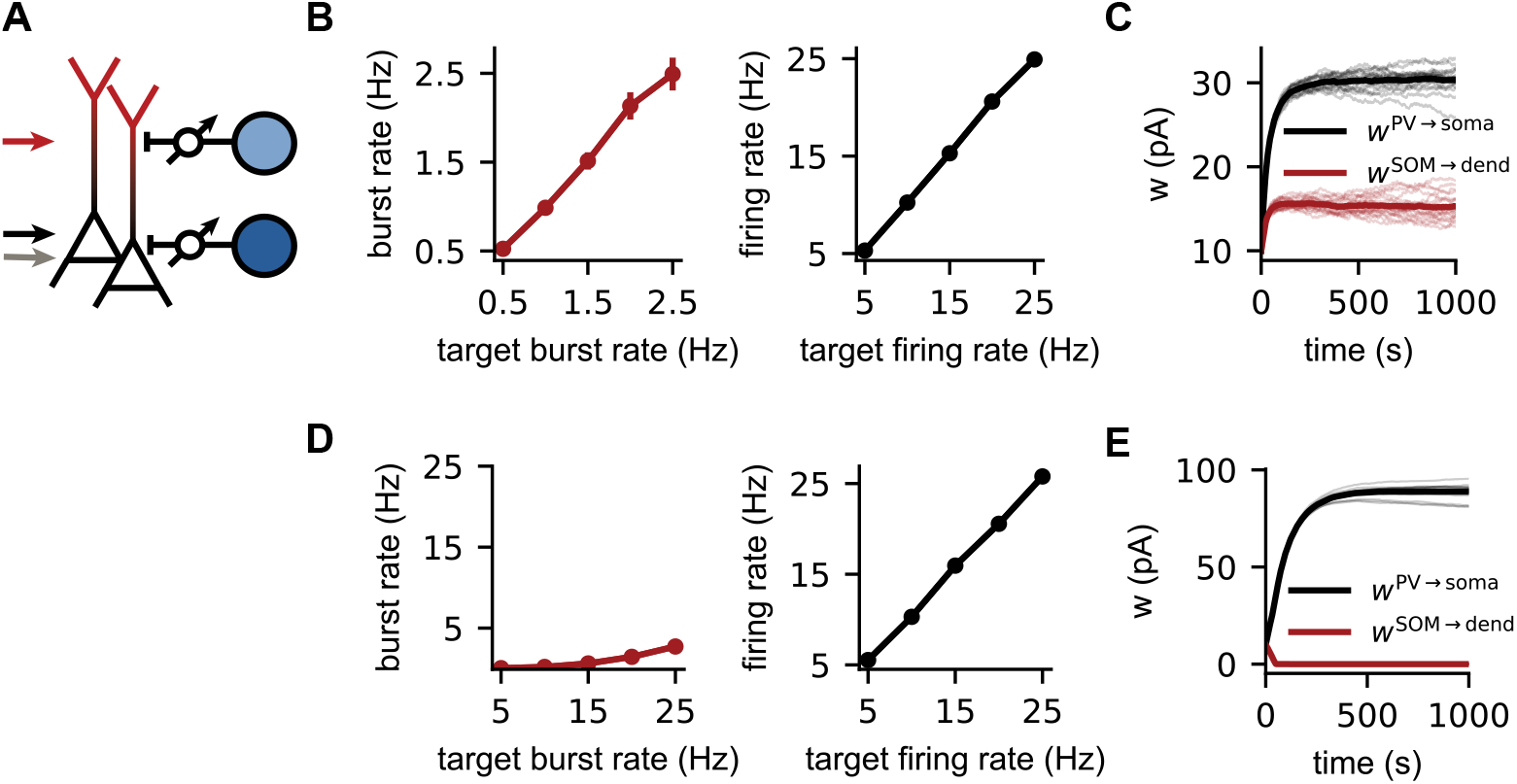
Simultaneous control of somatic and dendritic activity without and with competition between inhibitory plasticity rules. The somatic and dendritic activity of L5 PCs is under control of plastic inhibitory connections from PV (dark blue) and SOM (light blue) interneuron populations (see Fig 2. The somatic and dendritic compartments receive strong external inputs with moderate noisy background input. (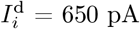, 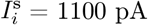, σ^d^ = σ^s^ = 100 pA). **(A, B)** No competition (target firing rate = 10 times target burst rate) versus **(C,D)** competition (target firing rate = target burst rate) between the target burst rate and target firing rate. **(A,C)** The burst and firing rate for different burst and firing target rates after learning the inhibitory weights **(B,D)**.

**Figure S4.**
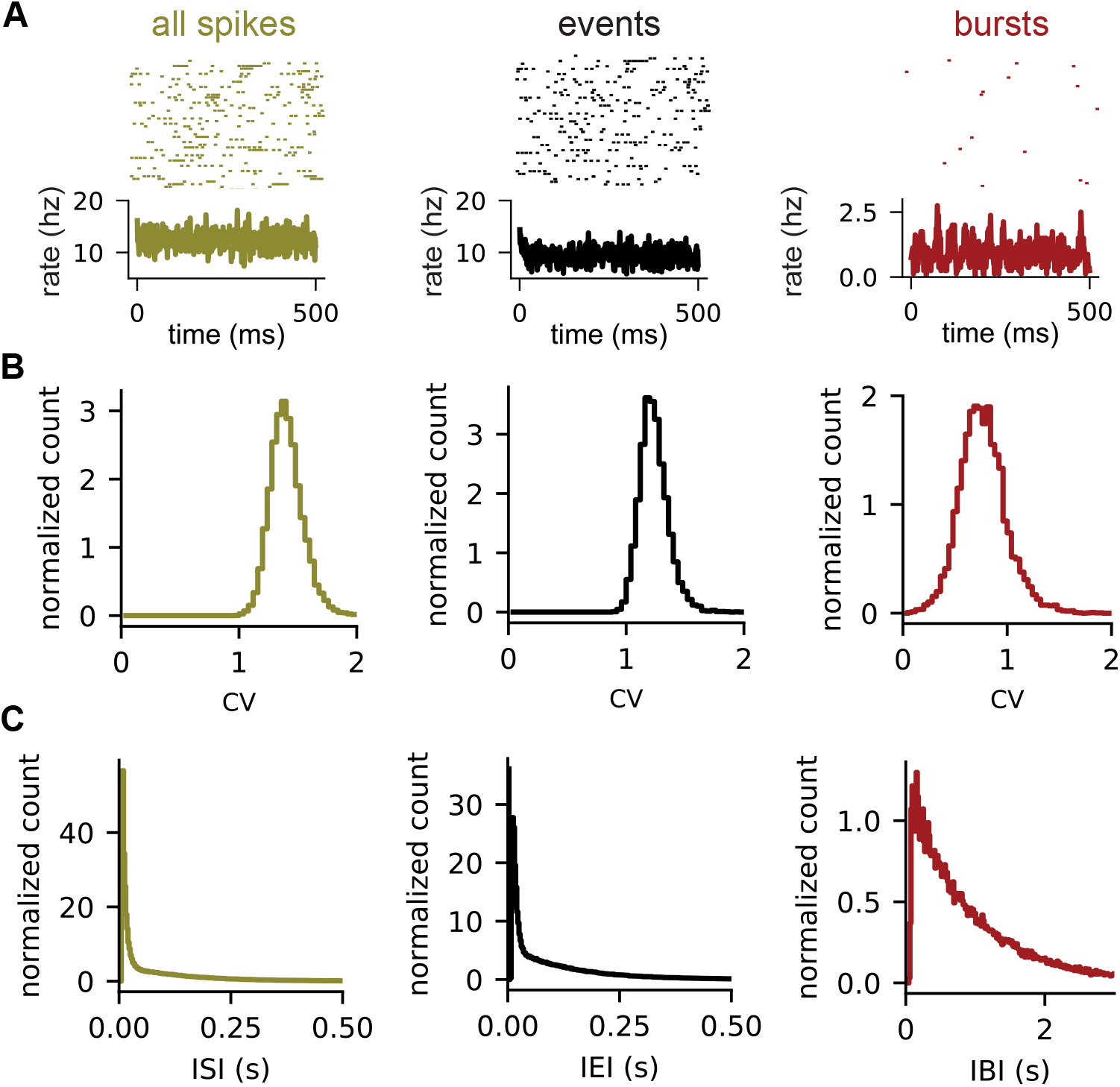
A doubly asynchronous irregular state for both spikes and bursts. The network configuration and stimulus condition are the same as for Fig 3 **(A)** (Top) Representative raster plots of all spikes, events and bursts of 50 neurons after learning the inhibitory weights. (Bottom) Histogram of all spikes, events or bursts of the entire PC population, normalized by the number of neurons (8000) and binsize (1 ms) to have units of rate. **(B)** The distribution of the coefficient of variation of the inter-spike intervals (CV ISI, yellow), inter-event intervals (CV IEI) and inter-burst intervals (CV IBI) after learning the inhibitory weights. **(C)** The distribution of the inter-spike intervals (ISI, yellow), inter-event intervals (IEI) and inter-burst intervals (IBI) after learning the inhibitory weights.

